# Direct and latent effects of ocean acidification on the transition of a sea urchin from planktonic larva to benthic juvenile

**DOI:** 10.1101/2021.12.10.471756

**Authors:** Narimane Dorey, Emanuela Butera, Nadjejda Espinel-Velasco, Sam Dupont

## Abstract

Ongoing ocean acidification (OA) is expected to affect marine organisms and ecosystems. While sea urchins can survive a wide range of pH, this comes at a high energetic cost, and early life stages are particularly vulnerable. Information on how OA affects transitions between life-history stages is scarce. We evaluated the direct and indirect effects of pH (pH_T_ 8.0, 7.6 and 7.2) on the development and transition between life-history stages of the sea urchin *Strongylocentrotus droebachiensis*, from fertilization to early juvenile. Continuous exposure to low pH negatively affected larval mortality and growth. At pH 7.2, formation of the rudiment (the primordial juvenile) was delayed by two days. Larvae raised at pH 8.0 and transferred to 7.2 after competency had mortality rates five to six times lower than those kept at 8.0, indicating that pH also has a direct effect on older, competent larvae. Latent effects were visible on the larvae raised at pH 7.6: they were more successful in settling (45%) and metamorphosing (30%) than larvae raised at 8.0 (17 and 1% respectively). These direct and indirect effects of OA on settlement and metamorphosis have important implications for population survival.

## Introduction

As a consequence of rising atmospheric anthropogenic carbon dioxide (CO_2_), ocean acidification (OA) is expected to decrease the average surface ocean pH by 0.2-0.4 units by 2100^1,2^. While open ocean pH values present small variability throughout the year, coastal systems are characterized by natural daily and seasonal pH variations^3^. For instance, the sampling site (Gullmarsfjord, Sweden) in the present study is characterized by a yearly variation of surface seawater pH between 8.6 and 7.6, while the average pH remains 8.1^4^. Given the intrinsic variable character of coastal ecosystems, it is expected that by 2100, the lowest pH experienced by organisms in these areas will exceed the projected average pH of 7.7 for open ocean^5^.

Lowered pH significantly impacts survival, development, physiology and growth in many benthic invertebrates, some of which play important roles as bioturbators or keystone species^6^. A review of 49 research articles on the effect of OA on sea urchins listed a majority (60%) of negative effects, most of them sub-lethal^7^ (see also^8^), although the observed effects appeared species, population and stage specific^9^. Early-life stages seem to be particularly sensitive as OA increases metabolic rates and decreases growth rates in sea urchin larvae (e.g. for *Strongylocentrotus droebachiensis* used in this study^4^). Acidified water also affects other physiological parameters in sea urchin larvae such as acid-base regulation ^e.g. 10^, feeding ^e.g. 11^ or gene expression related to immunity ^e^.^g^. ^12^. One current hypothesis is that exposure to lower pH increases the energy cost to maintain vital functions, including homeostasis^10^, leading to less energy available for growth.

Such energy limitations could induce latent shortcomings, in particular during life-stage transitions, which require major morphological and physiological changes and are often energy demanding (e.g. sea urchin^13^, barnacle^14^, abalone^15^). In sea urchins, embryos develop into planktotrophic pluteus larvae that eventually reach the ability to settle (i.e. competency, generally reached in 20-25 days post-fertilization at 10-12°C in *S. droebachiensis*^16,17^). A competent larva develops a large embryonic urchin inside its body, the rudiment. After finding suitable settlement substrate and attaching to it, full metamorphosis into a benthic juvenile can be achieved^18–20^. While these transitions are likely to be susceptible to OA, most studies have been restricted to a single stage. There is however growing interest in investigating the effect of OA on the transitions between at least two succeeding life-history stages in invertebrates whether in sea urchins^21,22^, sea stars^23,24^, oysters^25–27^, corals^28,29^ or slipper limpets^30^. In particular in sea urchins, recent studies^31–33^ have investigated how larval exposure to lowered pH influences the juvenile stages.

The aim of our study was to investigate whether pH affects the transition from a planktonic larva to a benthic juvenile in the green sea urchin *S. droebachiensis*. We tested three hypotheses in order to untangle direct and latent effects of pH (**Fig. 1**). The first hypothesis (H_1_) tested whether “*continuous exposure to low pH throughout the development has a negative effect on larval, newly settled and early juvenile development*”. We compared responses of individuals exposed to three pHs (8.0, 7.6 and 7.2) from post-fertilization (Dpf=0, Days post-fertilization) to post-settlement/juvenile stages (Dpf=40). The second hypothesis (H_2_), examined whether “*low pH experienced by competent larvae have a direct negative effect on the settlement and metamorphosis success*”. We then raised larvae at pH 8.0 until competency (Dpf 29) and examined the response when transferred to three pHs (8.0, 7.6 and 7.2) until post-settlement/juvenile stages (Dpf 40). The last hypothesis (H_3_) tested whether “*settling individuals and metamorphosed juveniles from larvae raised under low pH are impaired relative to the ones raised under pH 8.0*” (i.e. latent effect). For testing H_3_, we raised larvae at three different pHs (8.0, 7.6 and 7.2) until competency (Dpf 29) and measured the responses of individuals transferred to a pH of 8.0 until post-settlement/juvenile stages (Dpf 40).

**Figure 1.**
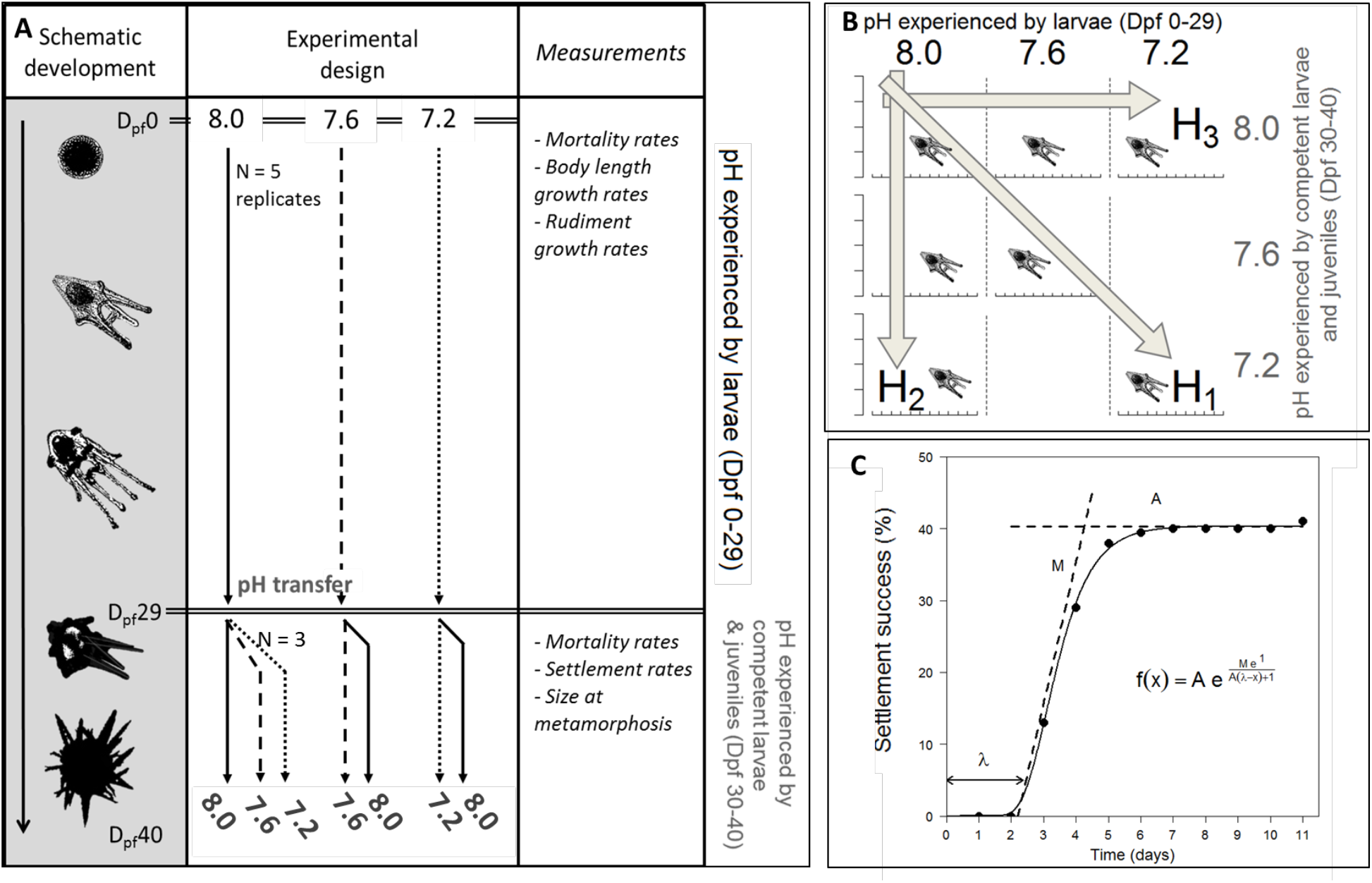
A. Timeline of the experiment. The experiment started (Day post-fertilization, Dpf 0) with cleaving embryos that were distributed into five replicate jars at three different pHs (8.0, 7.6 and 7.2). At Dpf 29, larvae from three of the five replicate jars were distributed individually into wells inside three 24-well plates per new pH condition (i.e. n=72 larvae for each treatment). **B. Experimental design**: Two parameters were tested: (i) pH experienced by larvae (Dpf 1 to 29) and (ii) pH experienced by stages from competent larvae to juveniles (Dpf 30-40). Seven combinations were tested allowing to test three different hypotheses: (H_1_), continuous exposure to low pH throughout the development has a negative effect on larval, competent, newly settled and early juvenile development; (H_2_), low pH experienced by competent larvae have a direct negative effect on the settlement and metamorphosis success; and, (H_3_), settling individuals and metamorphosed juveniles from larvae raised under low pH are impaired relative to the ones raised under pH 8.0 (i.e. latent effect). **C. The Gompertz model** used for settlement, with three descriptive parameters: maximum cumulated settlement (A; % larvae), maximum settlement rate (M; % larvae day-1) and initial latency period (λ; days from Dpf 30).

## Material and Methods

### (1) Adult collection and maintenance

The green sea urchin *S. droebachiensis* is a calcifying species widely distributed in boreal coastal ecosystems^34^. Adults *S. droebachiensis* were collected by divers in the Kattegat (Dröbak, Norway), transferred to the Sven Lovén Centre for Marine Sciences - Kristineberg facilities (Fiskebäckskil, Sweden) and maintained in 400-L basins with natural flowing seawater (seawater replacement: ≈1 L min^-1^), allowing a minimum volume of 10 L per individual. Sea urchins were kept at 12°C and fed *ad libitum* with *Ulva* spp. for four months prior to experiments.

### (2) Spawning and fertilization

In April 2012, spawning of the adults was induced by intra-coelomic injection of 1mL of 0.5 M KCl in filtered seawater (FSW). Eggs from four females were collected in FSW (≈9°C) while sperm from three males was collected dry and kept on ice. Sperm stock solution in FSW was subsequently added to an egg dilution to reach a final concentration of ≈1000 sperm mL^-1^, allowing a fertilization success above 95%. After fertilization (≈15 min), eggs were rinsed twice with FSW. Cleaving embryos (two-cell stage) were placed in 5-L culture jars filled with FSW at the target pH and at a density of ten embryos per mL in continuously aerated conditions.

### (3) General experimental design

An experiment was designed to test the impact of two different types of exposure to low pH (**Fig. 1A**): (i) pH experienced during the first phase of the larval development prior to the development of competency (well-formed rudiment; Dpf 0 to 29); and, (ii) pH experienced after the formation of the rudiment, settlement, metamorphosis and the juvenile stage. Three pHs were compared (8.0, 7.6 and 7.2) for a total of seven combinations of treatments (**Fig. 1A** and **1B**). This design allowed testing our three different hypotheses on direct and indirect effect of pH on transition from a planktonic larva to a benthic juvenile.

### (4) Larval incubation experiment (Dpf 0 to 29)

Larvae were cultured in glass jars until reaching competency at 29 days post-fertilization (Dpf 29) in three pH treatments (8.0, 7.6 and 7.2, **Fig. 1A**), and each treatment was replicated five times. Larval and later settler cultures were maintained at a temperature of 9.04 ± 0.25°C (mean ± sd) and in a salinity of ≈ 32.79 ± 0.25.

Target pHs in the culturing vessels (pH 7.2 and 7.6 ± 0.02 pH units) were achieved by bubbling pure CO_2_ in each bottle (pH computer AquaMedic). pH stability was visually assessed from the pH computer display at least twice a day. pH_NBS_, temperature and total alkalinity (TA) were measured in each jar twice a week. pH was monitored using a Metrohm (827 pH lab) electrode and adjusted from pH measurements on the total scale (pH_T_) using TRIS (Tris/HCl) and AMP (2-aminopyridine/HCl) buffer solutions with a salinity of 32 (provided by Unité d’Océanographie Chimique, Université de Liège, Belgium). Total alkalinity was assessed on filtered samples with a titration system (Titroline alpha plus, SI Analytics), following recommendations by Dickson et al.^35^. The carbonate system parameters (*p*CO_2_, Ω_Ca_ and Ω_Ar_) were calculated from measured temperature, pH_T_ and TA using the package *seacarb*^36^ in the software *R*^37^ with default values, following recommendations by Dickson et al.^35^ (see detailed results in supplementary information: **Table S1**).

From Dpf 6 to 25, larvae were fed daily with *Rhodomonas sp*. at a constant concentration of 150 μg C L^-1^ (3 000 – 6 000 cells mL^-1^ for diameters ranging between 6 and 8 μm). *Rhodomonas sp*. strains were provided by the Marine Algal Culture Centre of Gothenburg University (GUMACC) and maintained in B1 medium^38^ at 20°C under a 12:12 h light:dark cycle. Algal concentration and size were measured daily in every bottle using a Coulter counter (Elzone 5380, Micrometrics, Aachen, Germany) and concentration in each bottle was adjusted accordingly. At the chosen algal concentration (150 μg C L^-1^) and time of exposure (24 h, algae added daily), seawater pH has no impact on algal growth and survival^22^. As feeding can delay settlement and metamorphosis, feeding was stopped when a rudiment was visible (Dpf 25).

From Dpf 1 until 24, daily samples from each culture (2 x 10 mL) were fixed with 4% buffered paraformaldehyde in FSW and counted in order to estimate density. To account for developmental delay at low pH^39^, relative mortality rates (% larvae μm^-1^) were calculated for each replicate as the coefficient of the linear relationship between the relative density and body length rather than time. When the rates were non-significant (p-value>0.05), the mortality was considered null. At the four-arms stage (Dpf 10-20), larval morphology was scored as normal or abnormal after His et al.^40^ (i.e. very strong arm asymmetry, wrong arm orientation, deformities).

Every day, ten fixed larvae were randomly selected from each culture (Dpf 1-24) and photographed under a microscope (Leica with DFC295 camera). Body length (BL) and rudiment diameter were measured using the software ImageJ^41^. Larval growth rate was calculated for each replicate as the coefficient of the significant logarithmic relationship between BL and time. Two rudiment growth rates were calculated for each replicate as the coefficient of the significant linear relationship between rudiment diameter and (i) time post-fertilization (μm day^-1^) and (ii) BL (μm μm_BL_^-1^). Only rates calculated from significant relationships were used in subsequent analyses (p-value<0.05).

### (5) Settlement experiment (Dpf 30 to 40)

At Dpf 29, larvae showed signs of competency and presented visible, large (>100 μm) and opaque rudiments^42^. Competent larvae were transferred to settlement conditions for 11 days (i.e. until Dpf 40, **Fig. 1A**). A total of two-hundred and sixteen larvae were picked from three randomly selected larval cultures at pH 8.0, measured and distributed into 24-multiwell plates previously soaked in FSW for 24h (one larvae per 3 mL well; considered as replicate). The wells were filled with 3 mL of FSW pre-equilibrated at the target pH and temperature (≈9.0°C). Three plates were filled with FSW at pH 7.2, three with pH 7.6 and three with pH 8.0 (24 larvae x 3 plates x 3 pH treatments = 216 larvae). A similar process was used for larvae grown at 7.6 and 7.2, but larvae were only transferred to two pH treatments, either the same pH treatment or 8.0 (**Fig. 1A**; 24 larvae x 3 plates x 4 treatments = 288 larvae).

Each well was closed with parafilm and the plates were kept underwater in three 4-L aquaria filled with seawater equilibrated at the target pH (controlled by AquaMedic System as described above). The FSW in each well was replaced every 48 hours with water equilibrated at the same pH. Over this interval of time, the pH inside the wells was relatively stable and only changed by less than 0.17 pH units: from 8.00 to 8.06 ± 0.05 pH_T_ units; from 7.60 to 7.72 ± 0.12 pH_T_ units and from 7.20 to 7.37 ± 0.05 pH_T_ units (n=3 measurements of pooled samples from the seawater inside the wells). No food or settlement cues were added during this settlement period.

Every day, each individual was carefully followed through microscopy and scored as post-larva, newly settled post-larva (i.e. tightly attached to the inner surface of the wells), metamorphosed juvenile or dead. Each newly metamorphosed juvenile was photographed under a stereomicroscope (Leica MZ 16A with a DFC 420C camera) and the body diameter was measured. Relative mortality rate (% individuals day^-1^) was calculated as per the larval experiment. The cumulative settlement (in %) followed a Gompertz growth model and was analyzed as such using the package *grofit*^43^ in *R*. Three parameters were extracted from the model: maximum cumulated settlement (A; % larvae), maximum settlement rate (M; % larvae day^-1^) and initial latency period (λ; days; see **Fig. 1C**).

### (6) Data and statistical analyses

All the analyses were run with the software *R*^37^ applying a statistical significance threshold at α = 0.05. The normality of data distributions was checked with a Shapiro-Wilk test and homoscedasticity was tested using the Bartlett test. Results are given as mean ± SD.

For the larval experiment, culturing jars were true replicates (Dpf 0-29). In accordance with the data distribution, one-way ANOVA, Kruskall-Wallis (KW), Wilcoxon (W) or t-test (t) were used to test the effects of the pH treatments on the carbonate chemistry and biological parameters. Pairwise comparisons using t-tests with pooled SD with a Bonferroni adjustment were applied where significant effects were found.

For the settlement experiment, a generalized linear mixed effects model (GLMM, R package *nlme*^44^, function *lme*) was used to take into account a potential random effect of the plates. The effect of pH treatment, time (fixed factors) and plates (random factor) on relative mortality was tested. These GLMM were compared to GLMM without pH treatment or without time using ANOVA, to test for statistical significance of each of these factors in the original model. If models were significantly different (*p* < 0.05), model selection was done by comparing AIC (Akaike Information Criterion, higher quality model for lower AIC). A similar process was used to test for the statistical significance of the random factor: the initial GLMM was compared to a linear model without random factor (function *gls* from package *nlme*). For the proportion of settlement data, we log-transformed time before the analysis.

For the settlement experiment, we also tested the effect of pH on the three parameters (A, M and λ) extracted from the Gompertz models for each plate (ANOVA for the three Gompertz parameters, n=3 data points per pH treatment). In this second analysis, time was not included as a factor as it is already included in the Gompertz model. Plates are thus considered as the replication units (and not as a random factor), and A, M and λ are analyzed with one-way ANOVA (one fixed factor: pH treatment).

## Results

### (1) Carbonate chemistry conditions

During the *larval experiment* (Dpf 0 to 29), mean pH_T_ for each of the treatments was 8.00 ± 0.03 (*p*CO_2_ of 458 ± 41 μatm, Ω_ar_ = 1.73 ± 0.11 and Ω_ca_ = 2.74 ± 0.18), 7.60 ± 0.07 (*p*CO_2_ of 1262 ± 241 μatm, Ω_ar_ = 0.74 ± 0.11 and Ω_ca_ = 1.18 ± 0.18) and 7.24 ± 0.08 (*p*CO_2_ of 3027 ± 685 μatm, Ω_ar_ = 0.33 ± 0.05 and Ω_ca_ = 0.53 ± 0.08; see detailed results in **Table S1**). pH treatments were significantly different from one another (KW: χ^2^ = 105.9, df = 2 and *p* < 0.001) and pH did not vary between replicates within each pH treatment (KW: χ^2^ = 2.2, df = 4 and *p* = 0.70 for pH 7.2; χ^2^ = 4.5, df = 4 and *p* = 0.34 for 7.6 and χ^2^ = 7.4, df = 4 and *p* = 0.12 for 8.0). Temperature (9.04 ± 0.25°C) and TA (2321 ± 53 μmol kg^-1^) did not vary significantly between treatments (KW; Temperature: χ^2^ = 2.1, df = 2 and *p* = 0.35; TA: χ^2^ = 0.72, df = 2 and *p* = 0.70) nor replicates (KW; Temperature: χ^2^ = 21.1, df = 14 and *p* = 0.098; TA: χ^2^ = 11.5, df = 14 and *p* = 0.65).

### (2) Larval incubation experiment (Dpf 0 to 29)

During the larval incubation experiment, we followed the effect of continuously lowered pH on larval mortality and growth (H_1_).

Mean larval densities at Dpf 24 were 2.0 ± 1.2 larvae mL^-1^ at pH 7.2, 1.4 ± 1.5 larvae mL^-1^ at pH 7.6 and 2.5 ± 3.0 larvae mL^-1^ at pH 8.0. Larval densities in five cultures (two 7.2 replicates, one 7.6 replicate and two 8.0 replicates) were excluded from these calculations as they were too low to be recorded. Mortality rates (Dpf 0-24) were significantly higher at pH 7.2 than at pH 8.0 (0.22 ± 0.08 and 0.12 ± 0.03 % larvae μmBL^-1^ respectively), while rates at 7.6 were intermediary (0.17 ± 0.05% larvae umBL^-1^; ANOVA: F_2,12_ = 4.2, *p* = 0.040; post-hoc test; **Fig. 2A**). At the four-arms stage (Dpf 10-20), larval abnormality was significantly higher at 7.2 than in the two other treatments (ANOVA: F_2,12_ = 15.5, *p* = 0.005; post-hoc test). The proportion of abnormal larvae was 14 ± 18% for pH 8.0, 8 ± 12% for pH 7.6 and 26 ± 24% for pH 7.2.

**Figure 2.**
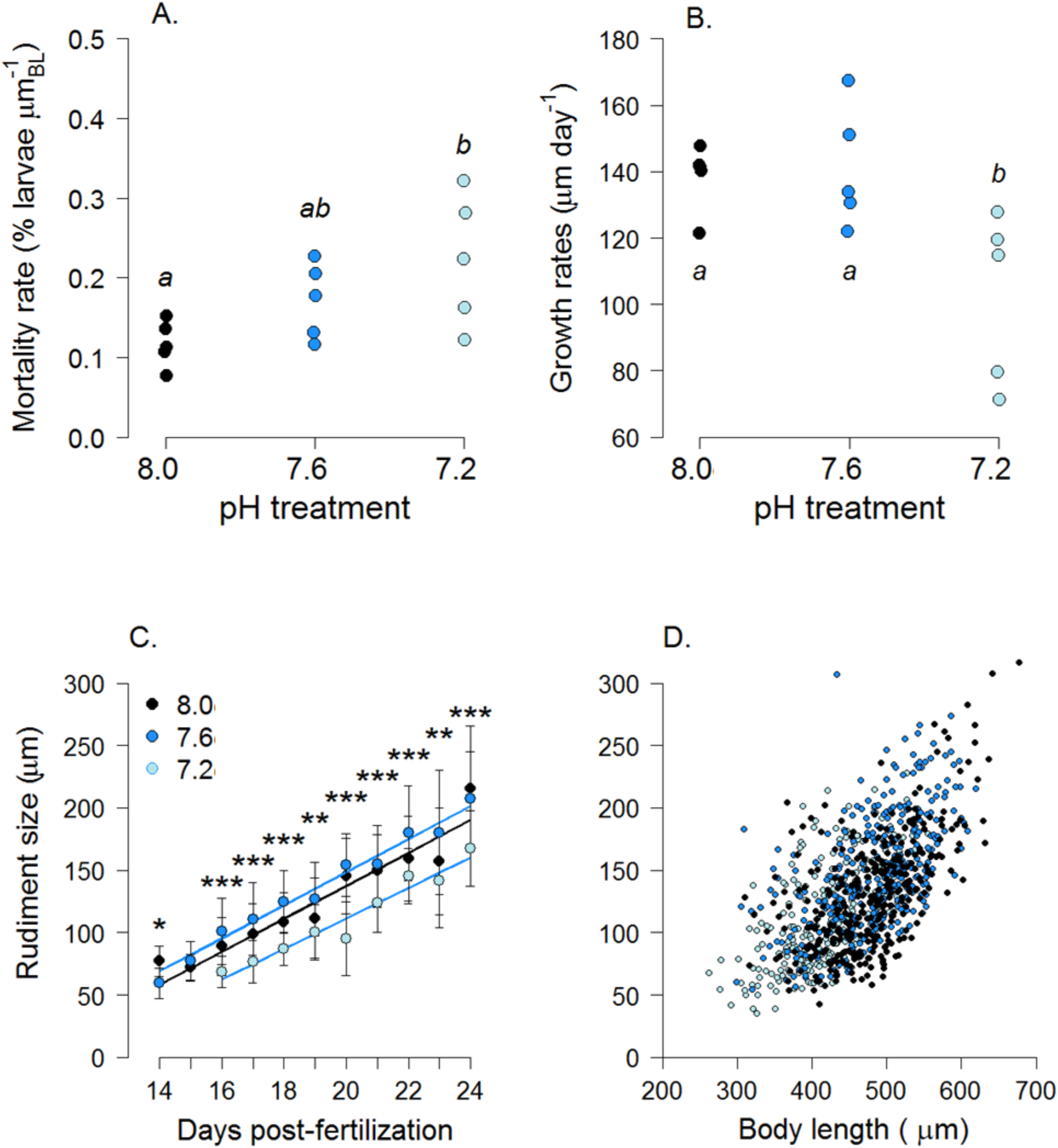
Effect of the pH treatments on the larval development (Dpf 1-29, H_1_): **A. mortality rates** (% larvae umBL^-1^), **B. body growth rates** (umBL day^-1^) and **C. rudiment size** (μm; mean±SD, vs. days post-fertilization); stars represent the significance of the pH effect (KW test; ** for *p*-value <0.01, *** for *p*-value <0.001; n=5 replicates per pH treatment). **D.** Rudiment size was significantly correlated to larval body size.

Larval body growth (**Fig. 2B**) was significantly lower in pH 7.2 (103 ± 25 μmBL log(day)^-1^) compared to pH 7.6 and 8.0 (141 ±18 and 139 ± 10 μmBL log(day)^-1^ respectively; ANOVA: F_2,12_ = 6.5, *p* = 0.012; post-hoc test). Rudiment growth was similar in all pH treatments (12 ± 4 μm log(day)^-1^; ANOVA: F_2,9_ = 0.3, *p* = 0.77). However, the rudiment appeared two days later in 7.2 and the size of the rudiment was always lower in this treatment (KW test on each day; **Fig. 2C**). Rudiment size was correlated to the body length (**Fig. 2D**, *p*<0.0001), albeit some dispersion (R^2^ = 39%), following the relationship *rudiment size* = *0.44 body length - 71.26*.

### (3) Settlement experiment (Dpf 30 to 40)

During the settlement experiment, we followed the effect of pH on competent larvae mortality, settlement and metamorphosis success, following three hypotheses H_1_, H_2_ and H_3_.

#### a. H_1_: Continuous exposure to low pH throughout the development has a negative effect on larval, competent, newly settled and early juvenile development

##### Mortality

For individuals maintained in the same pH treatment from larvae to settlement and metamorphosis, the pH treatment had no effect on competent larvae mortality (**Fig. 3A;** GLMM, models comparison: *p* = 0.21; see details in **Table S2**). Mean mortality rates ranged from 4.4 to 1.3 % per day (R^2^ > 86%).

**Figure 3.**
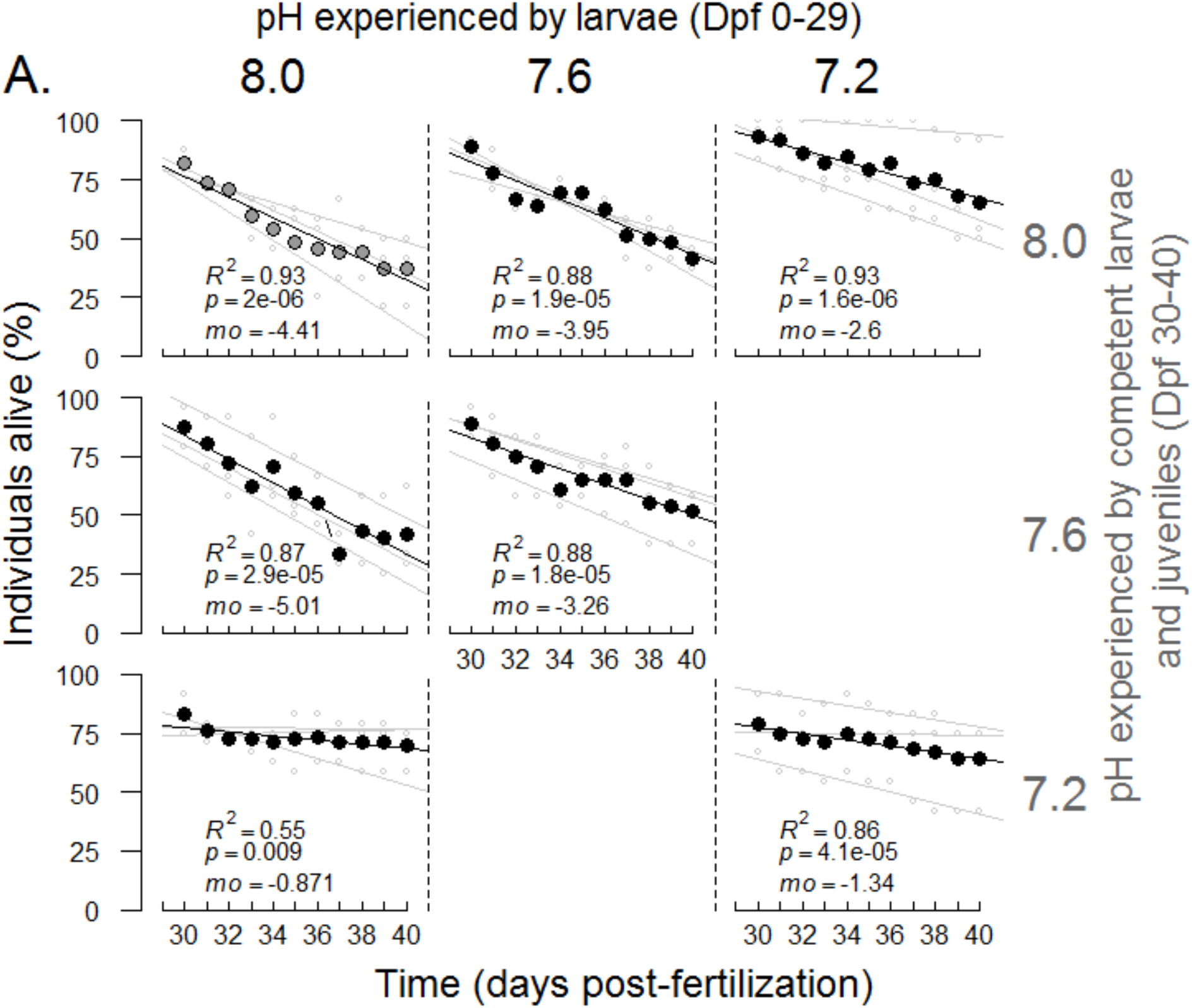
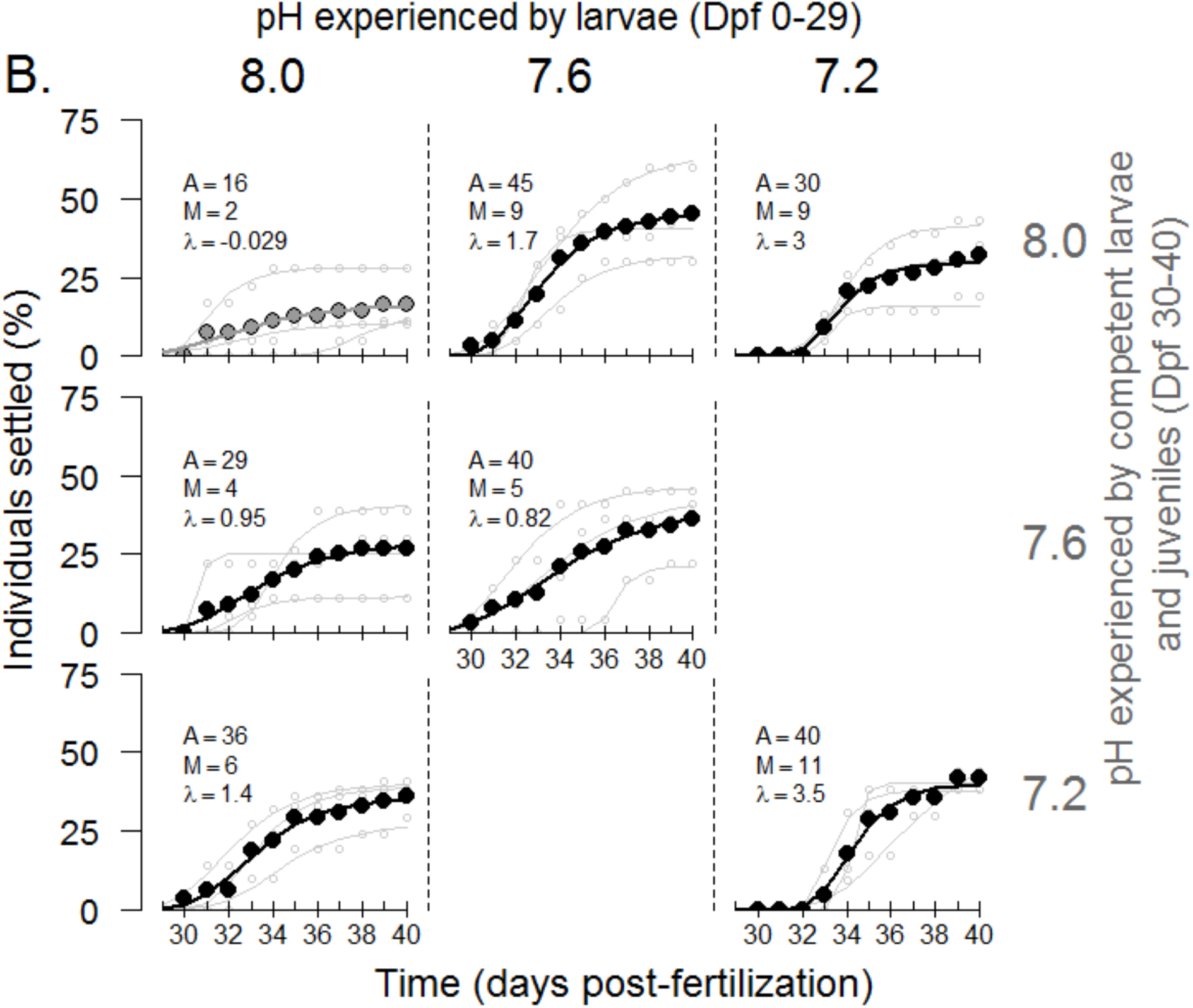
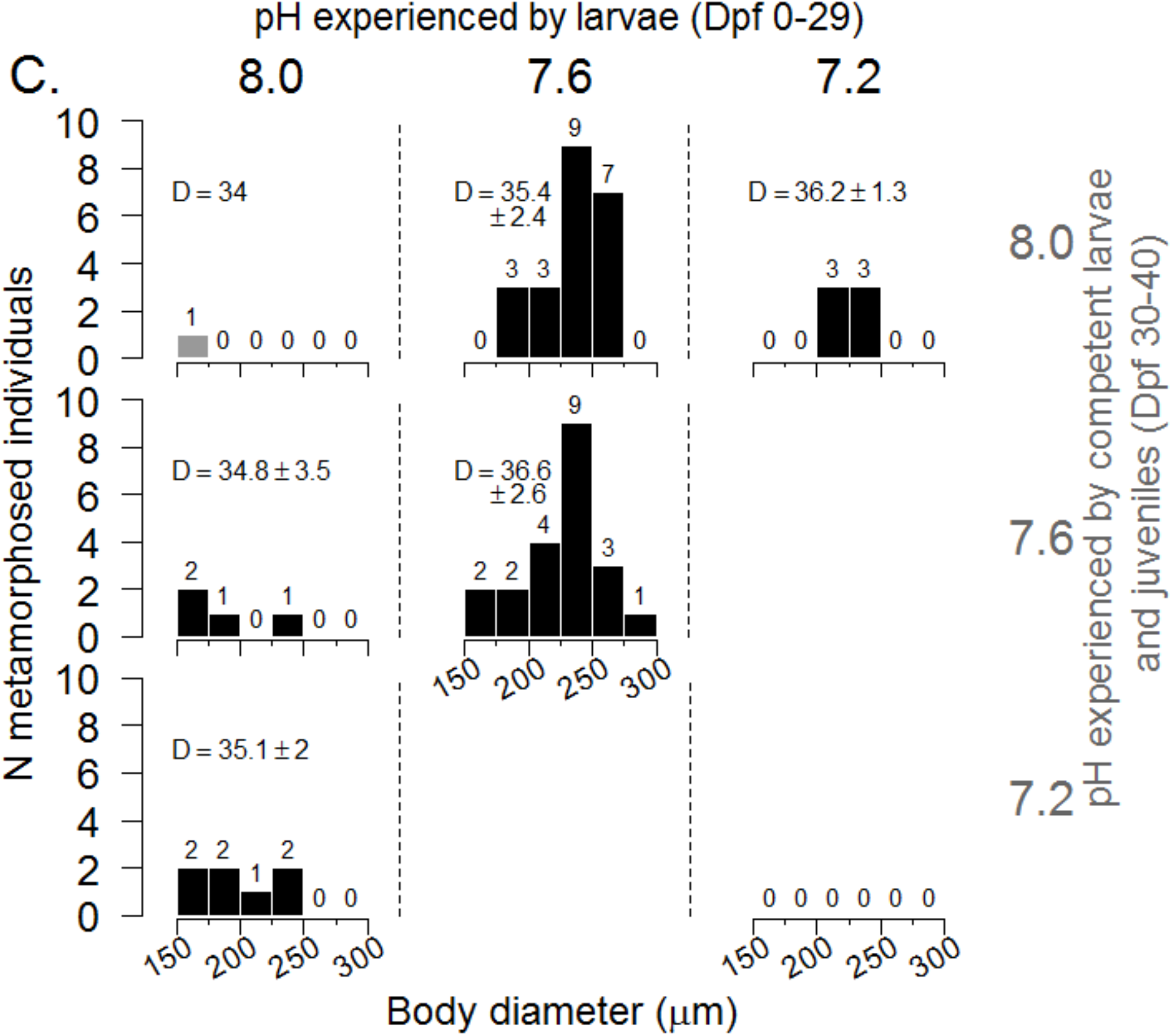
Effect of the combination of pH treatments on the settlement experiment (Dpf 29-40): At Dpf29, individuals from three pH (“*pH experienced by larvae*”, vertical pHs) were distributed in one of three pH treatments (horizontal pHs, see Fig. 1B). Full data points represent proportions calculated on the three well-plates combined (n=72), while light grey data points are calculated in each of the wellplates (n=24). **A. Mortality rates** (*mo*, % indiv. day^-1^) were calculated as the linear relationship between survival (individuals alive, %) and days. **B. Settlement success (%)** were fitted with Gompertz models (see **Fig. 1B**). A is the maximum cumulated settlement (% larvae), M is the maximum settlement rate (% larvae day^-1^) and λ is the initial latency period (days). Results of the models are detailed in **Table 1. C. Distribution of the newly metamorphosed individuals by body diameter at metamorphosis (μm)**. *D* represents the mean time (±SD) to metamorphosis (Dpf).

##### Settlement

pH treatment had no significant effect on the settlement dynamics (GLMM, models comparison: *p* = 0.22; details in **Table S2**). Similarly, none of the parameters of the settlement dynamics from the Gompertz model were significantly influenced by pH (**Table 1**). At the end of the settlement experiment, the average maximum cumulated settlement (A) was 34 ± 14 % (ANOVA, *p* = 0.057; **Fig. 3B**); average settlement rate (M) was 12 ± 9 % day^-1^ (*p* = 0.18); and the average initial latency period (λ) at the start of the settlement experiment was 2.7 ± 1.9 days (*p* = 0.80).

**Table 1:**
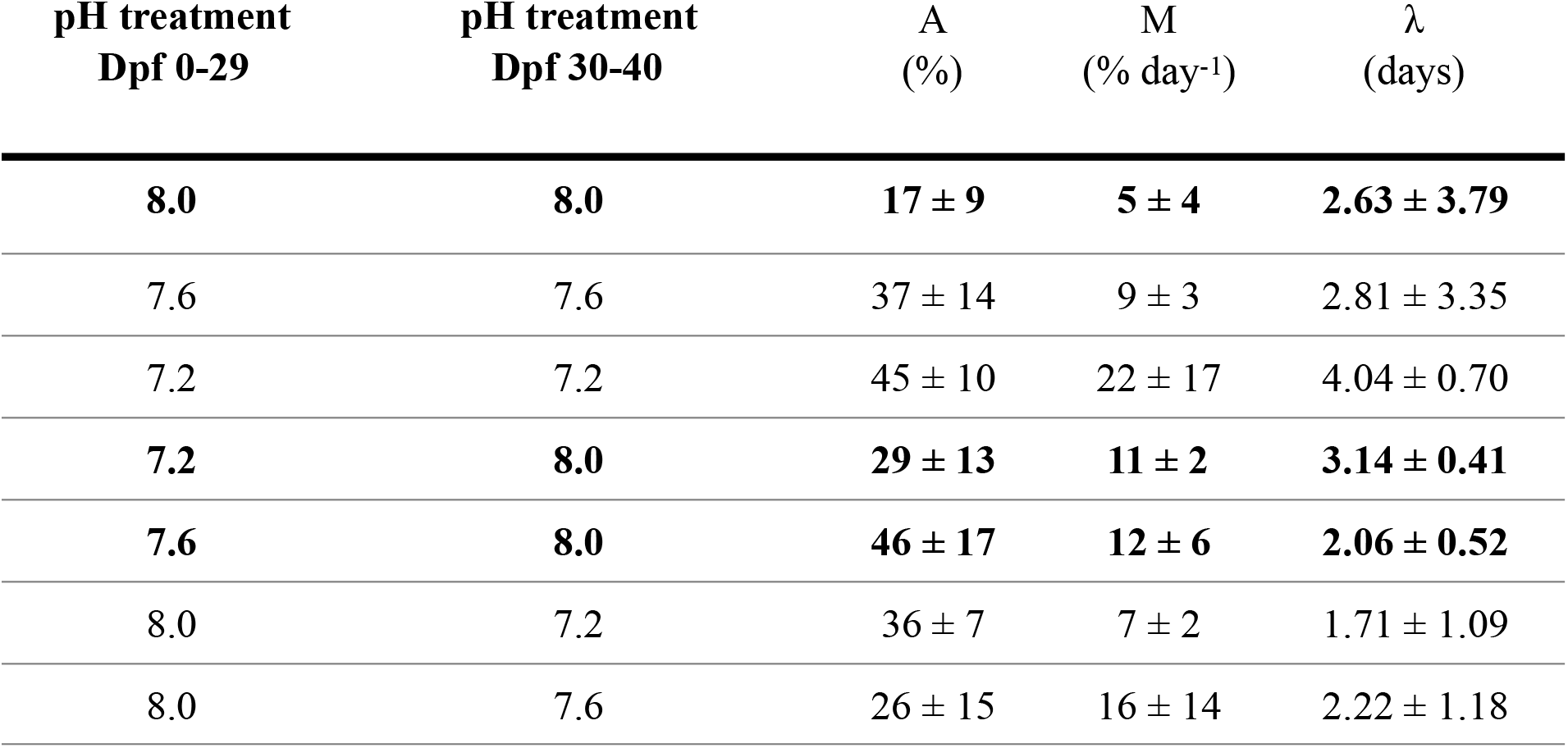
Parameters extracted from the Gompertz model by treatment (means ± SD, n=3 plates): maximum cumulated settlement (A; % larvae), maximum settlement rate (M; % larvae day-1) and initial latency period (λ; days from d30). The three treatments in bold were significantly different from each other.

| pH treatment<br>Dpf 0-29 | pH treatment<br>Dpf 30-40 | A<br>(%) | M<br>(% day<sup>-1</sup>) | $\lambda$<br>(days) |
| --- | --- | --- | --- | --- |
| **8.0** | **8.0** | **17 ± 9** | **5 ± 4** | **2.63 ± 3.79** |
| 7.6 | 7.6 | 37 ± 14 | 9 ± 3 | 2.81 ± 3.35 |
| 7.2 | 7.2 | 45 ± 10 | 22 ± 17 | 4.04 ± 0.70 |
| **7.2** | **8.0** | **29 ± 13** | **11 ± 2** | **3.14 ± 0.41** |
| **7.6** | **8.0** | **46 ± 17** | **12 ± 6** | **2.06 ± 0.52** |
| 8.0 | 7.2 | 36 ± 7 | 7 ± 2 | 1.71 ± 1.09 |
| 8.0 | 7.6 | 26 ± 15 | 16 ± 14 | 2.22 ± 1.18 |

#### b. H_2_: Low pH experienced by competent larvae have a direct negative effect on the settlement and metamorphosis success

##### Mortality

Larvae raised at pH 8.0 and transferred to 7.2 at Dpf 30 had mortality rates five to six times lower than the ones kept at pH 8.0 or 7.6 (~ 0.8, 4.2 and 5 % day^-1^ for pH 7.2, 7.6 and 8.0, respectively, **Fig. 3A**). This direct effect of pH during the settlement period was significant (GLMM, *p* = 0.0495, **Table S2**).

##### Settlement

pH experienced during the larval phase and by competent larvae did not significantly influence their settlement dynamic (GLMM, *p* = 0.20, **Table S2**) or the settlement parameters extracted from the Gompertz model (ANOVA: *p* = 0.20 for A, *p* = 0.32 for M and *p* = 0.90 for λ; **Table 1** and **Fig. 3B**).

#### c. H_3_: settling individuals and metamorphosed juveniles from larvae raised under low pH are impaired relative to the ones raised under pH 8.0 (“latent effect”)

##### Mortality

The pH in which larvae were raised had a significant effect on the mortality of competent larvae maintained at pH 8.0 (GLMM, *p* **=** 0.021, **Table S2**).The average mortality rates for larvae raised at pH 7.6 (~ 3.8 % day^-1^, R^2^ = 88%) were similar to those raised at pH 8.0 (~ 4.2 % day^-1^) but higher than those raised at 7.2 (~ 2.5 % day^-1^, R^2^ = 93%; **Fig. 3A, Fig. S1**).

##### Settlement

The pH in which larvae were raised had a significant effect on the settlement dynamic (GLMM, *p* = 0.044, **Table S2**). For example, (**Fig. 3B**) nearly half of the individuals settled when larvae were raised at pH 7.6 (maximum settlement, A=46 ± 17 %) when only 29 ± 13 % of the larvae raised at pH 7.2 did and 17 ± 9 % for pH 8.0. However, pH had no significant effect on any of the settlement parameters extracted from the Gompertz model (ANOVA: *p* = 0.10 for A, *p* = 0.17 for M and *p* = 0.84 for λ, **Table 1, Fig. S2**).

### (4) Metamorphosis (Dpf 30 to 40)

*H_1_*: Individual did not metamorphose after being raised continuously at pH 8.0 and 7.2, whereas 30% did at pH 7.6 (**Fig. 3C**, **Fig. S3**). As a consequence, it was not possible to test the effect of pH on the size of the newly metamorphosed individuals or the timing of metamorphosis. *H_2_:* Few individuals (6-10%) metamorphosed after being transferred from pH 8.0 to a lower pH (pH 7.6 or 7.2). *H_3_:* Larvae transferred from pH 7.6 to pH 8.0 had the highest metamorphosis success (31%), while 8% (n=6) metamorphosed after being transferred from pH 7.2 to pH 8.0.

## Discussion

We tested three different hypotheses related to the direct and indirect effect of seawater pH experienced during larval development, settlement, metamorphosis and juvenile stages.

### H_1_: Continuous exposure to low pH throughout the development has a negative effect on larval, newly settled and early juvenile development

#### Larval development

Larvae exposed to pH 7.2 (Dpf 0-29) presented higher mortality and abnormality rates, as well as lower growth rates than their counterparts at pH 7.6 and 8.0. In addition, rudiments developed later at the lowest pH 7.2. Growth delay and abnormalities have typically been observed in sea urchin larvae under acidified conditions (see reviews^7,8^). In *S. droebachiensis* larvae, metabolism (respiration^4^ and feeding^11^) increased due to exposure to lowered pH. This increased metabolism - possibly due to increased/additional costs of certain physiological functions (e.g. pH regulation and digestion^10,11,45^) – could increase maintenance costs and thus reduce scope for growth^39,46^.

A change in energy allocation could also explain the significant delay in formation of the rudiment as well as the increased mortality and abnormality at pH 7.2 during the pre-settlement period. pH-induced changes in morphology (abnormalities, asymmetry) such as those observed here could be a consequence of a disruption of one or more molecular mechanisms involved in development^12^. Building the rudiment requires a considerable energy investment from the larvae ^e.g. 47^, and lowering the pH can be expected to negatively impact the subsequent transitions. Interestingly, rudiment growth rate (**Fig. 2C**) and size relative to the same larval body length was similar in all three pH conditions (**Fig. 2D**). The smaller rudiment observed at 7.2 at a given time can hence be attributed to the larval growth delay rather than to an additional effect of larval energy restriction on the rudiment. This result suggests that the building of the rudiment is a priority of the larval energy investment. Accordingly, Vaïtilingon et al.^48^ reached similar conclusions as they observed a faster growth of the rudiment than of the larval body (2.3-fold) in fed competent *P. lividus* larvae.

#### Settlement

When competent larvae (Dpf 30-40) were exposed at settlement to the same pH as the treatment in which they had grown, pH did not have a statistically significant effect on the mortality (**Fig. 3A**) or the settlement parameters (cumulated settlement, rates of settlement or latency period, **Fig. 3B**). Comparing our results to other published studies is difficult (e.g. ^49,50^) as different protocols were used (e.g. no inducer used in our experiment) and we defined settlement as substrate attachment while others defined settlement as the installation of settlers (no difference between settled competent larvae or juveniles). Nevertheless, and in contrast to our results, García et al.^49^ observed that, in the sea urchin *P. lividus*, lowering pH induced a delay in the settlement of competent larvae by eight days at pH 7.7 (vs. 8.1), and no settlement was found at pH 7.4. Howver, García et al.^51^ also found that settlement in the sea urchin *P. lividus* was unaffected at reduced pH (7.7 vs. 8.1) when competent larvae were presented with a suitable algal substrate.

#### Metamorphosis

In our study, only one out of 72 larvae metamorphosed into juvenile when raised at pH 8.0 and none at pH 7.2. Although we observed relatively high settlement at pH 7.2 (40%), it did not result in any juveniles. This could suggest that, for individuals grown at pH 7.2 from early stages, a full metamorphosis might be too costly to achieve. However, around 30% of the competent larvae raised at pH 7.6 metamorphosed (**Fig. 3C**). An early settlement behavior observed at 7.6 – together with increased metamorphosis success – can be interpreted in the light of the “desperate larvae hypothesis” proposed by Marshall and Keough^52^. This hypothesis postulates that a larva in unfavorable conditions (e.g. low food concentration or, for our study, the energetically challenging low pH conditions) would settle earlier, as an alternative to a risky or costly planktonic life. Although this strategy allows escape from unfavorable planktonic conditions, it limits the probability of finding an adequate substrate for juvenile growth, hence diminishing the chances of survival after metamorphosis. Under this assumption, larvae raised at pH 8.0 would still have enough energy reserve to delay their metamorphosis and wait for a more suitable substrate. As biologically active environments that are favorable for juvenile urchins are characterized by larger variability in pH, an alternative hypothesis would be that a mild decrease in pH (e.g. 7.6 vs. 8.0) could play the role of an inducer. Because of the low number of newly metamorphosed individuals in control conditions, we were unable to test the hypothesis that lower pH decreased settlers’ size (see results from Wangensteen et al.^50^).

### H_2_: Low pH experienced by competent larvae have a direct negative effect on the settlement and metamorphosis success

For larvae raised in pH 8.0 until competency (Dpf 29), mortality rates were five times lower for individuals transferred to pH 7.2 as compared to those transferred to pH 8.0 or 7.6, indicating that pH also has a direct effect on older, competent larvae. Our observation on the absence of direct effects of lowered pH on settlement is not isolated: settlement rates of *Centrostephanus rodgersii* were unaffected when competent larvae (Dpf 40) were transferred to lowered pH^31^ (7.6 and 7.8 vs. 8.1; see also studies on *Evechinus chloroticus*^53^ and *Psedechinus huttoni*^33^). In the present study, metamorphosis was higher when competent larvae were transferred to lower pH as compared to pH 8.0 (**Fig. 3C**). This observation supports the possibility that competent larvae could preferentially settle and metamorphose when encountering an environment with lower pH. Sea urchin larvae settle preferably on biologically active settlement substrates such as crustose coralline algae^54,55^. Given their photosynthesis/respiration activities, crustose coralline algae change the pH of the boundary layers leading pH variability and lower pHs than the surrounding waters^33^. pH might potentially be used as a proxy by competent larvae to identify suitable habitats. The effect of lower pH could also be indirect through a stress associated with increased energy costs and leading to metamorphosis (desperate larvae hypothesis).

### H_3_: settling individuals and metamorphosed juveniles from larvae raised under low pH are impaired relative to the ones raised under pH 8.0 (“latent effect”)

For larvae grown at reduced pH until competency (Dpf 29) and subsequently transferred to pH 8.0, average mortality rates were significantly influenced by the pH experienced during early larval development. The lowest daily mortality rates were found for pH 7.2 (2.5% vs. 4% in pH 8.0). This could be explained by a latent effect as at Dpf 30, competent larvae raised at 7.2 were smaller and had smaller rudiments. To understand if reduced mortality at low pH stems from the developmental delay ^e^.^g^.^56^, future studies will need to monitor the energy reserves of competent larvae.

A latent effect was also observed for the settlement dynamics. The maximum settlement was higher for larvae raised at pH 7.6 (46%) as compared to the other two treatments (29% and 17% for pH 7.2 and 8.0, respectively). This could suggest that pH 7.6 may be the optimal pH for the larval development of this species. The importance to consider the present natural variability, extreme low pH in particular, to define species sensitivity has been highlighted by Vargas et al.^57^. *S. droebachiensis* spawn in spring during plankton blooms and are naturally exposed to a large pH variability ranging from 8.6 to 7.6^4^. As a consequence, a pH 7.6 would be within the species tolerance level and regularly experienced by the larval stages.

While we observed “positive” latent effects, the literature principally reports negative latent postsettlement effects of lowered pH on early-post settlement growth in invertebrates (see review^58^). For example, when reared at low pH (7.8) until metamorphosis and transferred to control pH conditions (8.1), oysters performed poorer (decrease in juvenile growth rates by 26%) than individuals raised at 8.1 all along^26^, an effect that persisted for at least four months after metamorphosis^27^. Coral recruits decreased their calcification and growth when larvae had previously been reared at low pH^29^. While we found latent effects that increased settlement success, experiments on the same species showed that prior exposure to pH 7.6 had a negative latent effect on the post-settlement survival and growth^22^. This could indicate that earlier settlement at pH 7.6 is not necessarily a “positive” latent effect for juvenile survival, with low pH having a direct impact on early juveniles test and spines^31,33^.

In conclusion, our results highlight pH as an important modulator of the transition between larval and juvenile stages. Larvae raised within the present range of pH environmental variability show some plasticity in their settlement and metamorphosis pattern while larvae raised at pH 7.2 never achieved metamorphosis. This confirms the presence of a physiological tipping point below pH 7.6 for this species^4,46^. Survival and effective recruitment of juveniles into the adult population has been seldom investigated in the context of OA and remains a black box today in the predictions of future population changes. This study highlights the potential of OA to modify the recruitment into adult populations through slower growth rates and decrease of the metamorphosis success at extreme low pH. Ocean acidification might lead to recruitment in an inhospitable environment that - in the long term - might modify urchin distributions^59^ and induce effects at the ecosystem level^60^.

## Acknowledgements

SD is funded by the CeMEB and supported by a Linnaeus-grant from the Swedish Research Councils VR and Formas. This work was performed within the Linnaeus Centre for Marine Evolutionary Biology, CeMEB (http://www.cemeb.science.gu.se/).

