## Supplementary Tables and Figures for "Direct and latent effects of ocean acidification on the transition of a sea urchin from planktonic larva to benthic juvenile"

**<u>SUPPLEMENTARY TABLES AND FIGURES FROM THE MANUSCRIPT:</u>**

### SUPPLEMENTARY TABLES AND FIGURES

**Table S1:** Seawater carbonate chemistry parameters during the *larval experiment* (Dpf 0-29) presented by pH treatment and replicate (R). Two seawater parameters from the carbonate system (total-scale pH: $pH_T$ and total alkalinity: TA) and temperature (T) were measured in each replicate twice a week during the larval experiment and used to calculate $CO_2$ partial pressure ($pCO_2$) as well as the aragonite and calcite saturation states (respectively Ωar and Ωca), for a salinity of 32.5 using the package *seacarb* for R (Lavigne & Gattuso 2011). All the values are expressed as mean ± SD. Note that the seawater was always under-saturated in the *larval-pH* 7.3 treatments, but only under-saturated with respect to aragonite in the *larval-pH* 7.7 treatments.

| **pH** | R | $pH_T$ | T (°C) | TA ($mmol.kg^{-1}$) | $pCO_2$ (µatm) | Ωar | Ωca |
|---|---|---|---|---|---|---|---|
| 7.2 | A | 7.24 ± 0.06 | 8.90 ± 0.25 | 2.31 ± 0.03 | 2933 ± 432 | 0.33 ±0.05 | 0.53 ±0.07 |
| | B | 7.25 ± 0.07 | 9.04 ± 0.22 | 2.34 ± 0.06 | 2928 ± 515 | 0.34 ±0.05 | 0.55 ±0.08 |
| | C | 7.23 ± 0.06 | 9.11 ± 0.25 | 2.29 ± 0.02 | 3045 ± 441 | 0.32 ±0.04 | 0.51 ±0.07 |
| | D | 7.24 ± 0.13 | 8.99 ± 0.31 | 2.37 ± 0.17 | 3177 ± 1309 | 0.35 ±0.08 | 0.55 ±0.12 |
| | E | 7.23 ± 0.06 | 9.03 ± 0.25 | 2.32 ± 0.02 | 3053 ± 455 | 0.32 ±0.04 | 0.51 ±0.06 |
| 7.6 | A | 7.62 ± 0.07 | 8.96 ± 0.21 | 2.32 ± 0.03 | 1191 ± 193 | 0.77 ±0.10 | 1.23 ±0.16 |
| | B | 7.55 ± 0.10 | 9.13 ± 0.23 | 2.33 ± 0.02 | 1435 ± 339 | 0.68 ±0.14 | 1.08 ±0.23 |
| | C | 7.63 ± 0.05 | 9.01 ± 0.20 | 2.31 ± 0.04 | 1157 ± 147 | 0.79 ±0.09 | 1.25 ±0.15 |
| | D | 7.61 ± 0.05 | 9.13 ± 0.23 | 2.31 ± 0.03 | 1208 ± 149 | 0.76 ±0.07 | 1.20 ±0.11 |
| | E | 7.59 ± 0.08 | 9.00 ± 0.23 | 2.33 ± 0.03 | 1318 ± 261 | 0.72 ±0.12 | 1.14 ±0.19 |
| 8.0 | A | 7.98 ± 0.03 | 9.30 ± 0.31 | 2.32 ± 0.04 | 486 ± 39 | 1.66 ±0.11 | 2.63 ±0.17 |
| | B | 8.01 ± 0.05 | 9.09 ± 0.31 | 2.32 ± 0.02 | 448 ± 51 | 1.76 ±0.15 | 2.79 ±0.24 |
| | C | 8.02 ± 0.04 | 8.93 ± 0.18 | 2.33 ± 0.03 | 436 ± 42 | 1.80 ±0.11 | 2.85 ±0.18 |
| | D | 8.00 ± 0.03 | 9.13 ± 0.19 | 2.31 ± 0.03 | 466 ± 34 | 1.70 ±0.09 | 2.69 ±0.14 |
| | E | 8.00 ± 0.02 | 8.91 ± 0.21 | 2.32 ± 0.03 | 452 ± 28 | 1.74 ±0.06 | 2.76 ±0.09 |

**Table S2:** Statistical results of the GLMM for post-competency **A.** mortality and **B.** settlement. A GLMM including the effect of pH treatment, time (fixed factors) and plates (random factor) was compared to a GLMM without pH treatment using ANOVA and the statistical results are given in this table (p-value for model comparison, AIC for each GLMM, equation for the best model and the significance of the effects of time and plates).

**A. <u>Mortality</u>**

| Hypothesis tested[†] | p-value for model comparison,<br>- Models AIC | *Equations for the best-fit model** | Best-fit model variables |
|---|---|---|---|
| H1 | $p = 0.21$<br>- with pH: 723<br>- without pH: 722 | *1) individuals alive =*<br>$81.9 - 3.0\ Time + R_{plate}$ | - Time: significant fixed effect ($p < 0.0001$)<br>- $R_{plate}$: significant random effect ($p < 0.0001$), distribution $N(0, 7.6^2)$ |
| H2 | $\boldsymbol{p = 0.0495}$<br>- with pH: 766<br>- without pH: 768 | *1) individuals alive in 8.0 → 7.2 =*<br>$93.6 - 3.4\ Time + R_{plate}$<br>*2) individuals alive in 8.0 → 7.6 =*<br>$79.4 - 3.4\ Time + R_{plate}$<br>*3) individuals alive in 8.0 =*<br>$75.0 - 3.4\ Time + R_{plate}$ | - Time: significant fixed effect ($p < 0.0001$)<br>- $R_{plate}$: significant random effect ($p < 0.0001$), distribution $N(0, 10^2)$ |
| H3 | $\boldsymbol{p = 0.021}$<br>- with pH: 717<br>- without pH: 721 | *1) individuals alive in 7.2 → 8.0 =*<br>$102.0 - 3.7\ Time + R_{plate}$<br>*2) individuals alive in 7.6 → 8.0 =*<br>$84.67 - 3.7\ Time + R_{plate}$<br>*3) individuals alive in 8.0 =*<br>$76.3 - 3.7\ Time + R_{plate}$ | - Time: significant fixed effect ($p < 0.0001$)<br>- $R_{plate}$: significant random effect ($p < 0.0001$), distribution $N(0, 7.5^2)$ |

*For the equations, **pH treatments** are referred to as *"larvae raised at pHx (Dpf 0-29) → competent larvae and juveniles raised at pHy (Dpf 30-40)"* (e.g. "8.0 → 7.2" for larvae raised at 8.0 and transferred to 7.2 after 29 Dpf) or simply e.g. "8.0" when larvae experienced only one pH treatment all along the experiment. When no treatment is mentioned, the pH treatment had no significant effect. †**Hypotheses**: in $H_1$ we compared the responses of larvae raised at 8.0, 7.6 or 7.2 from Dpf 0 to 40; in $H_2$ we compared treatments in which larvae were grown at 8.0 between Dpf 0 and 29; in $H_3$ ("latent effect") we compared treatments in which larvae were transferred to pH 8.0 at Dpf 29.

**B. <u>Settlement</u>**

| Hypothesis tested† | p-value for model comparison,<br>- Models AIC | *Equations for the best-fit model** | Best-fit model variables |
|---|---|---|---|
| H1 | $p = 0.22$<br>- with pH: -189<br>- without pH: -190 | 1) *individuals settled =*<br>*-0.046 + 0.143 log(Time) +* $R_{plate}$ | - Time: significant fixed effect ($p < 0.0001$)<br>- $R_{plate}$: significant random effect ($p < 0.0001$), distribution $N(0, 0.08^2)$ |
| H2 | $p = 0.20$<br>- with pH: -240<br>- without pH: -241 | 1) *individuals settled =*<br>*-0.011 + 0.115 log(Time) +* $R_{plate}$ | - Time: significant fixed effect ($p < 0.0001$)<br>- $R_{plate}$: significant random effect ($p < 0.0001$), distribution $N(0, 0.06^2)$ |
| H3 | $\mathbf{p = 0.044}$<br>- with pH: -200<br>- without pH: -198 | 1) *individuals settled in 7.2→8.0 =*<br>*-0.054- 0.145 log(Time) +* $R_{plate}$<br>2) *individuals settled in 7.6→8.0 =*<br>*0.113 - 0.145 log(Time) +* $R_{plate}$<br>3) *individuals settled in 8.0 =*<br>*-0.066 - 0.145 log(Time) +* $R_{plate}$ | - Time: significant fixed effect ($p < 0.0001$)<br>- $R_{plate}$: significant random effect ($p < 0.0001$), distribution $N(0, 0.07^2)$ |

**Figure S1**: **A. Mortality rates of competent larvae from Dpf 29 to 40** (*mo*, % indiv. day$^{-1}$) were calculated as the **B.** linear relationship between survival (individuals alive, %) and days.

Legend: 8.0c, 7.6c and 7.2c represent continuous pH ($H_1$), 8.0-7.6 and 8.0-7.2 represent a transfer from pH 8.0 to lower pH ($H_2$) and 7.6-8.0 and 7.2-8.0 the test for latent effects ($H_3$). Each circle represents the mean mortality calculated for one plate (n=24 individuals, random factor). The influence of the pH treatment was significant for $H_2$ and $H_3$ (see results in **Table S2**).

**A.**

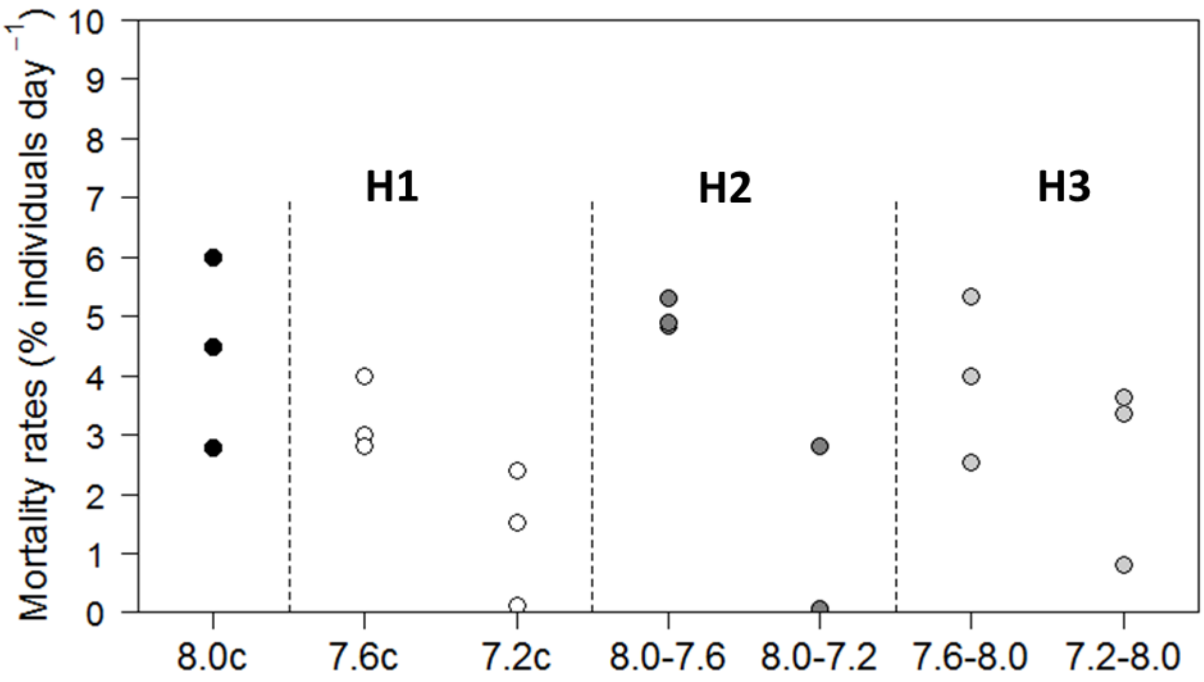

53 **B.**

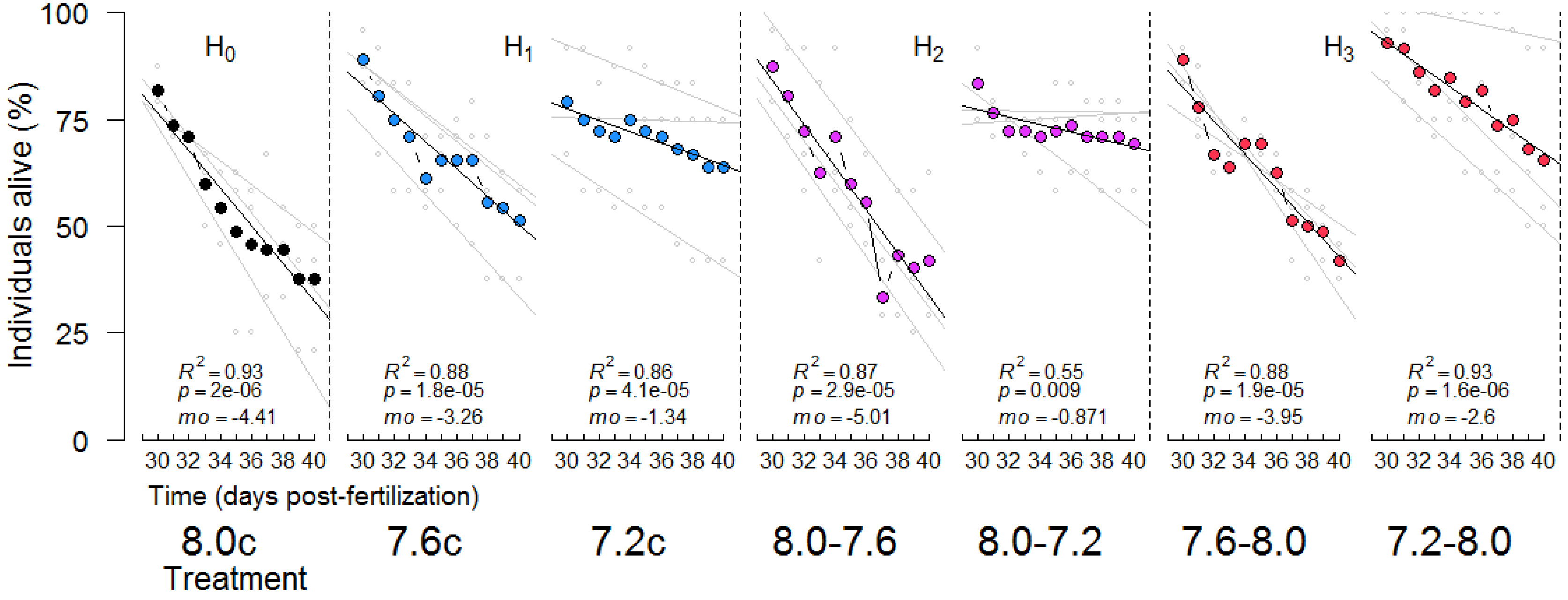
Individuals alive (%)
100
75
50
25
0
H0
H1
H2
H3
R2 = 0.93
p = 2e-06
mo = -4.41
R2 = 0.88
p = 1.8e-05
mo = -3.26
R2 = 0.86
p = 4.1e-05
mo = -1.34
R2 = 0.87
p = 2.9e-05
mo = -5.01
R2 = 0.55
p = 0.009
mo = -0.871
R2 = 0.88
p = 1.9e-05
mo = -3.95
R2 = 0.93
p = 1.6e-06
mo = -2.6
30 32 34 36 38 40
Time (days post-fertilization)
8.0c
7.6c
7.2c
8.0-7.6
8.0-7.2
7.6-8.0
7.2-8.0
Treatment

54

**Figure S2**: **Settlement success (%)** were fitted with Gompertz models. A is the maximum cumulated settlement (% larvae), M is the maximum settlement rate (% larvae $day^{-1}$) and $\lambda$ is the initial latency period (days). Results of the models are detailed in **Table 1**. Full data points represent proportions calculated on the three well-plates combined (n=72), while light grey data points are calculated in each of the well-plates (n=24).

Legend: 8.0c, 7.6c and 7.2c represent continuous pH ($H_1$), 8.0-7.6 and 8.0-7.2 represent a transfer from pH 8.0 to lower pH ($H_2$) and 7.6-8.0 and 7.2-8.0 the test for latent effects ($H_3$). The influence of the pH treatment was significant for $H_3$ (see results in **Table S2**).

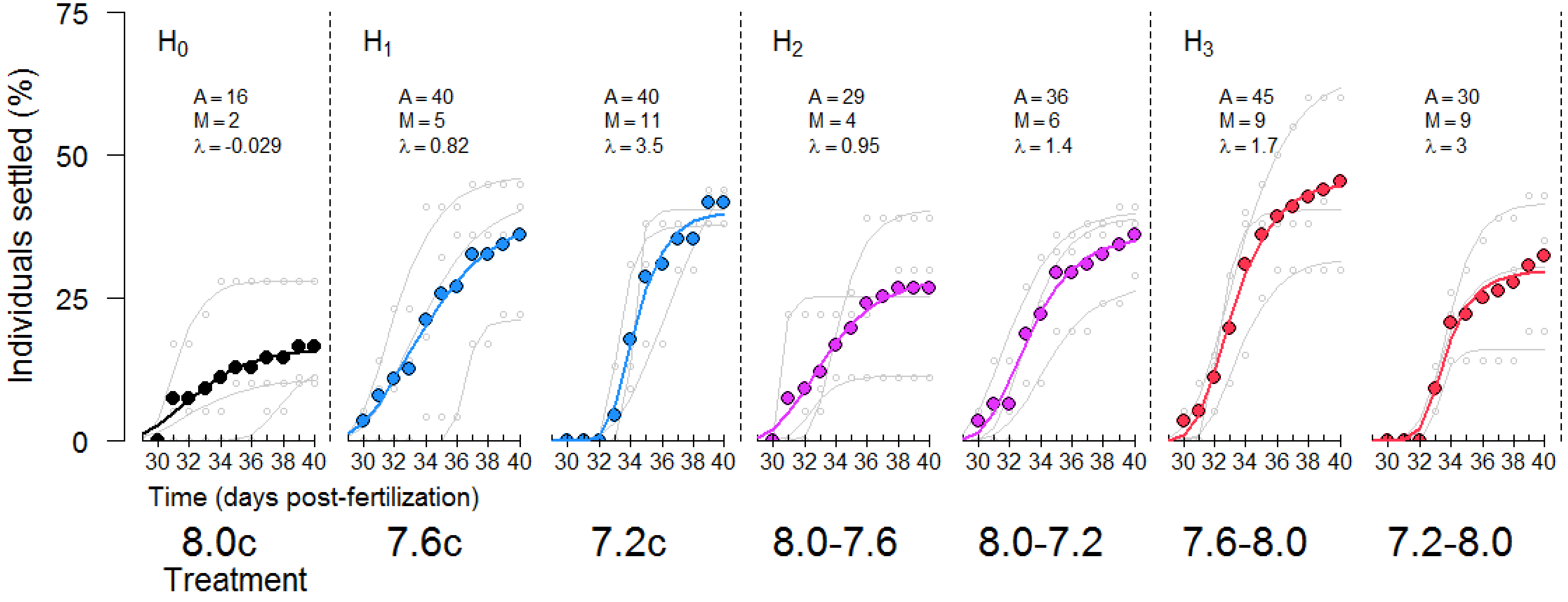

**Figure S3**: **Distribution of the newly metamorphosed individuals by body diameter at metamorphosis (µm)**. *D* represents the mean time (±SD) to metamorphosis (Dpf).

Legend: 8.0c, 7.6c and 7.2c represent continuous pH ($H_1$), 8.0-7.6 and 8.0-7.2 represent a transfer from pH 8.0 to lower pH ($H_2$) and 7.6-8.0 and 7.2-8.0 the test for latent effects ($H_3$). The influence of the pH treatment was not statistically tested.

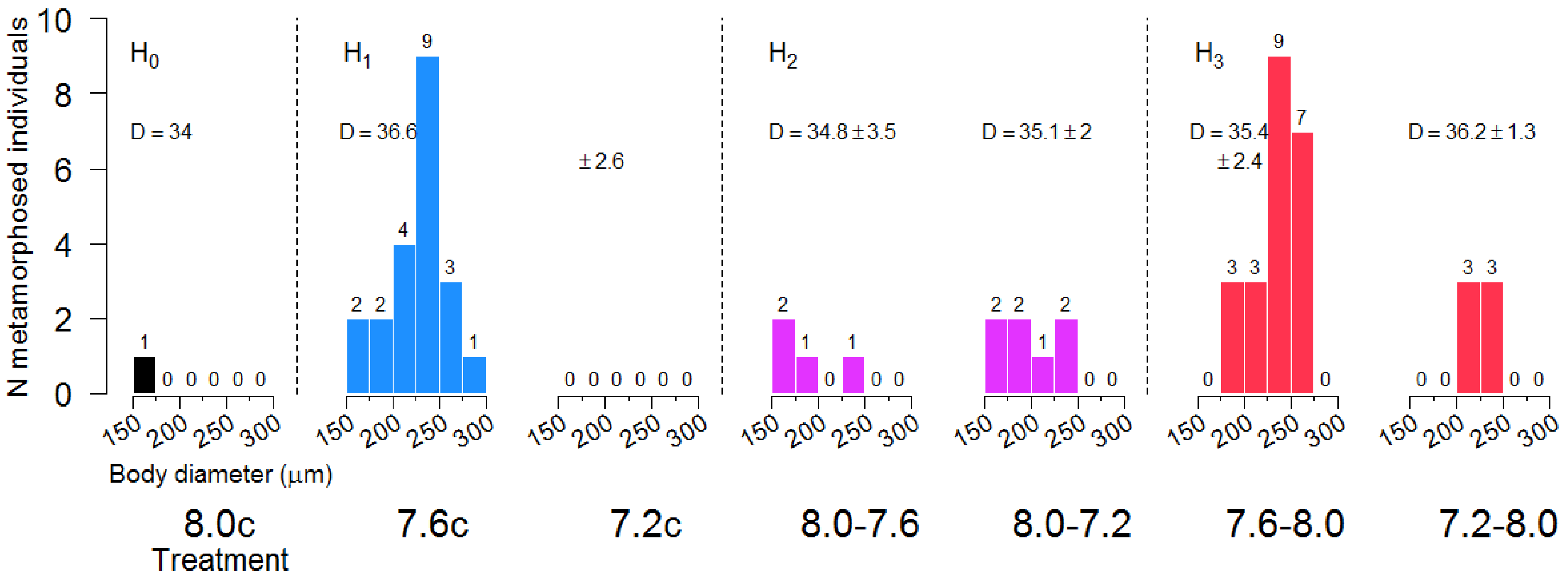
